# Structural basis for recognition of anti-migraine drug lasmiditan by the serotonin receptor 5-HT_1F_–G protein complex

**DOI:** 10.1101/2021.05.29.446083

**Authors:** Sijie Huang, Peiyu Xu, Yangxia Tan, Chongzhao You, Yumu Zhang, Yi Jiang, H. Eric Xu

## Abstract

Migraine headache has become global pandemics and is the number one reason of work day loss. The most common drugs for anti-migraine are the triptan class of drugs that are agonists for serotonin receptors 5-HT_1B_ and 5-HT_1D._ However, these drugs have side effects related to vasoconstriction that could have fatal consequences of ischemic heart disease and myocardial infarction. Lasmiditan is a new generation of anti-migraine drug that selectively binds to the serotonin receptor 5-HT_1F_ due to its advantage over the tripan class of anti-migraine drugs. Here we report the cryo-EM structure of the 5-HT_1F_ in complex with Lasmiditan and the inhibitory G protein heterotrimer. The structure reveals the mechanism of 5-HT_1F_-selective activation by Lasmiditan and provides a template for rational design of anti-migraine drugs.

---

The serotonin 5-HT_1_ receptor subtypes, including 5-HT_1A_, 5-HT_1B_, 5-HT_1D_, 5-HT_1e_, and 5-HT_1F_, are G protein-coupled receptors (GPCRs) that respond to the endogenous neurotransmitter serotonin and couple preferentially to the G_i/o_ family of G proteins^1^. Drugs targeting 5-HT_1_ receptors are used to treat migraine, depression, and schizophrenia^2^. Clinical use of traditional anti-migraine drugs, triptans, caused side effects arising from therapeutic vasoconstrictive actions when targeting 5-HT_1B/1D_ receptors^3^. The requirement of new anti-migraine drugs lacking vasoconstrictive effects led to the development of lasmiditan, a highly selective 5-HT_1F_ receptor agonist with minimizing on-target side effects^4^.

Migraine is one of the most common diseases worldwide and, importantly, a major cause of lost work productivity^3^. The selective 5-HT_1B/D_ agonists, triptans, are currently a first-line acute treatment of moderate-to-severe migraine attacks. Triptans bind mostly to 5-HT_1B/D_ receptors within cerebral blood vessels, leading to vasoconstriction. Unfortunately, a large percentage of patients are not satisfied with current acute migraine treatments, because 5-HT_1B/D_ receptors are also present on coronary and limb arteries and triptans may cause acute coronary syndromes in patients with or without cardiovascular disease^3,5^.

Lasmiditan, a potent and selective agonist for the 5-HT_1F_ receptor, has recently been approved for acute migraine^6^. Lasmiditan lacks vasoconstriction effects and may be a safer and more effective option for patients refractory to treatment with triptans and for patients with cardiovascular disease^6^. Lasmiditan has a pyridinyl-piperidine scaffold, which is structurally different from the indole derivatives of triptans. In addition, Lasmiditan is able to penetrate the blood-brain barrier to act on receptor located in the brain, thus enhancing its action on receptor sites in central nervous system (CNS)^6^. To better understand the structural basis of lasmiditan for the 5-HT_1F_ selectivity and activation, we determined the structure of the 5-HT_1F_ in complex with lasmiditan and G_i1_ at a resolution of 3.4 Å by single-particle cryo-EM. The structure reveals the mechanism of 5-HT_1F_-selective activation and provides a template for the rational design of anti-migraine drugs.

For single-particle cryo-EM structural studies, we prepared the lasmiditan-bound 5-HT_1F_–G_i_ complex, which were met with technical challenges of low expression levels and unstable formation of the receptor-G protein complex. Despite these difficulties and after many attempts, we were able to prepare homogenous sample for cryo-EM analysis. The structure was determined at a global resolution of 3.4 Å (Supplementary information, Fig. S1). The lasmiditan-bound 5-HT_1F_–G_i_ complex EM density maps are sufficiently clear to define the position of the 5-HT_1F_ receptor, the G_i_ heterotrimer, scFv16, and the bound ligand lasmiditan. The overall structure of 5-HT_1F_ consists of a canonical transmembrane domain (TMD) of seven transmembrane helices (TM1-7), a short intracellular loop 2 (ICL2) helix, and an amphipathic helix H8 (Fig. 1a, b). The active 5-HT_1F_ receptor shares a similar overall conformation with other active 5-HT_1_ receptors^7^, while a complete backbone structure for ECL2 is visible, which is partly missing in other 5-HT_1_ structures due to the flexibility. The cryo-EM map includes well-defined features for amino acids forming the agonist-binding pocket and clear density for lasmiditan in 5-HT_1F_ (Fig. 1b). We found that negatively charged amino acids in the ligand binding pocket of 5-HT_1F_ are primarily responsible for the affinity of lasmiditan (Fig. 1c, d). In the orthosteric binding pocket (OBP), the primary amine on methylpiperidine group of lasmiditan forms a canonical charge interaction with D103^3×32^ of 5-HT_1F_ (Fig. 1d), which simultaneously forms a hydrogen bond with Y337^7×42^, supporting a stable interaction between the ligand and the receptor (Fig. 1d). Mutational studies showed these residues are critical for lasmiditan binding (Supplementary information, Fig. S2). The interactions from D^3×32^ of the receptor to the primary amine of agonists and the supportive Y^7×42^ are conserved in aminergic GPCRs^8^. In addition, the methylpiperidine group of lasmiditan forms hydrophobic interactions with F309^6×51^ in TM6 of 5-HT_1F_ (Fig. 1d), mutation in F309^6×51^ cause a nearly 100-fold reduction of lasmiditan affinity (Supplementary information, Fig. S2). Meanwhile, the aromatic pyridine scaffold of lasmiditan is sandwiched between I104^3×33^ and F310^6×52^, forming a hydrophobic interaction core (Fig. 1d). F310^6×52^A mutation simultaneously eliminated 5-HT_1F_-G protein coupling signals and lasmiditan affinity, and I104^3×33^ A mutation also cause a nearly 60-fold reduction of lasmiditan affinity, suggesting that these hydrophobic interactions are crucial for lasmiditan-induced 5-HT_1F_ activation (Supplementary information, Fig. S2). I^3×33^ of 5-HT_1F_ is 3.5 Å away from the aromatic ring of lasmiditan, which provides a stronger hydrophobic interaction than V^3×33^ of 5-HT_1A_ and M^3×33^ of 5-HT_1E_. In the extended binding pocket (EBP), the trifluorobenzene group of lasmiditan forms additional hydrophobic interactions with I174^ECL2^ and P158^4×60^, and forms hydrogen bonds with residue E313^6×55,^ N317^6×59^, T182^5×40^, and H176^ECL2^ of 5-HT_1F_ (Fig. 1d). These structural observations are also confirmed by mutation experiments (Supplementary information, Fig. S2).

**Fig. 1.**
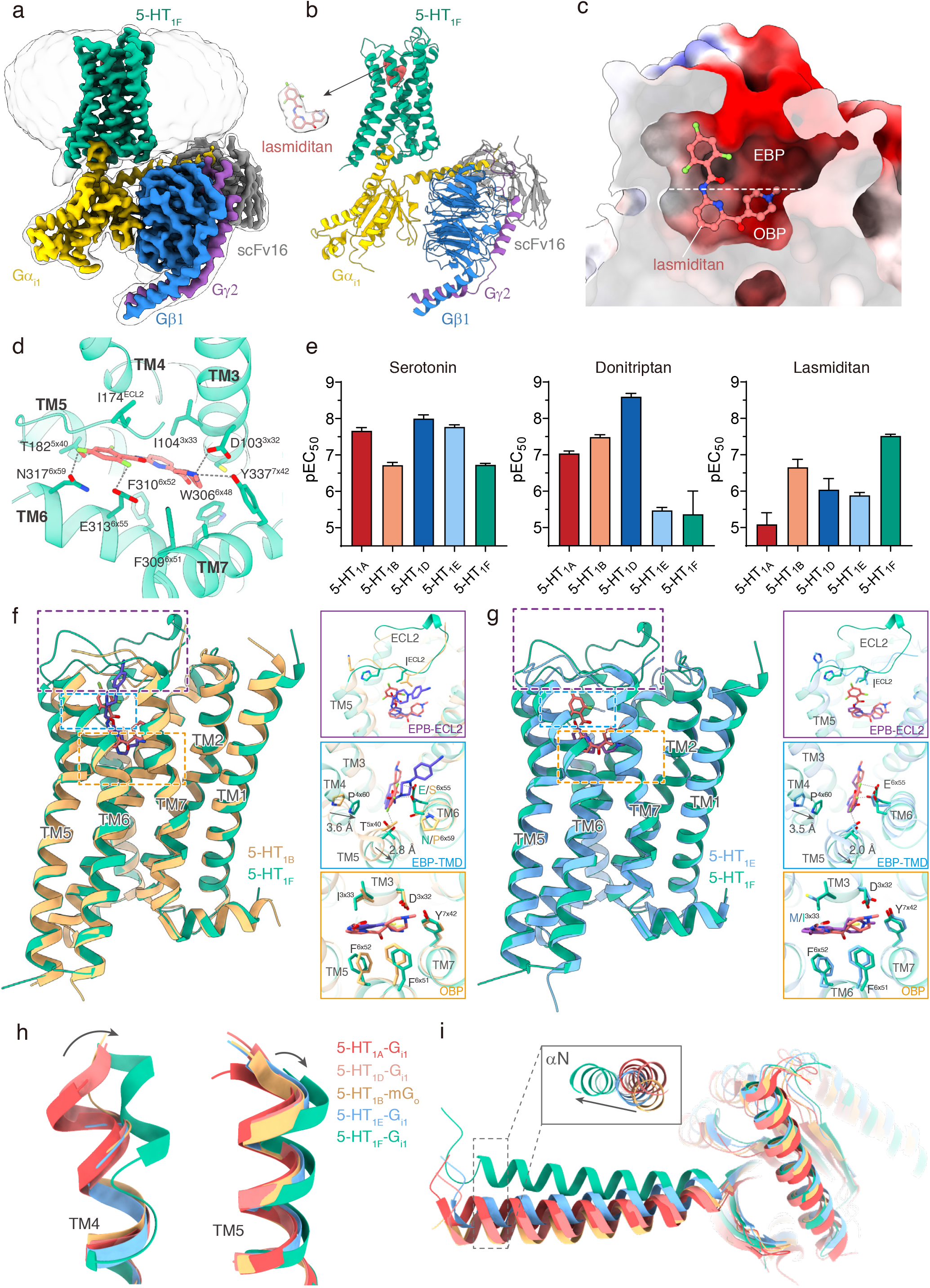
Structure of lasmiditan–5-HT_1F_–G_i1_ complex. **a** Cryo-EM map of the 5-HT_1F_–G_i_ complex. **b** Structural model of the 5-HT_1F_–G_i_ complex. The ligand model is shown on left side of the complex with surrounding density map. **c** Electrostatic surface representation of lasmiditan binding pocket of 5-HT_1F_. **d** The binding mode of lasmiditan in the ligand binding pocket of 5-HT_1F_. **e** G_i_ recruitment assay using NanoBiT for wild type 5-HT_1A_, 5-HT_1B_,5-HT_1D_, 5-HT_1E_, and 5-HT_1F_ induced by serotonin, donitriptan and lasmiditan. **f** Structural comparison of lasmiditan-bound 5-HT_1F_ with donitriptan-bound 5-HT_1B_ (PDB code: 6G79). **g** Structural comparison of lasmiditan-bound 5-HT_1F_ with BRL54443-bound 5-HT_1E_ (PDB code: 7E33). **h** Structure comparison focus on extracellular end of TM4 (left) and TM5 (right) among 5-HT_1A_ (red, PDB code: 7E2Y), 5-HT_1B_ (tan, PDB code: 6G79), 5-HT_1D_ (yellow, PDB code: 7E32), 5-HT_1E_ (blue, PDB code: 7E33) and 5-HT_1F_ (green). **i** Comparison of the G_α_ conformation among the structures of G_i/o_ coupled 5-HT_1A_ (red, PDB code: 7E2Y), 5-HT_1B_ (tan, PDB code: 6G79), 5-HT_1D_ (yellow, PDB code: 7E32), 5-HT_1E_ (blue, PDB code: 7E33) and 5-HT_1F_ (green).

Lasmiditan is a new generation 5-HT receptor agonist with high affinity and selectivity for 5-HT_1F_ (Ki = 2 nM) over other serotonin receptors (Ki > 500 nM)^4^, and this selectivity is also confirmed by our NanoBiT G-protein recruitment assays (Fig. 1e; Supplementary information, Fig. S3a). Comparison of the structure of 5-HT_1F_ bound to lasmiditan with other 5-HT_1_ structures^7,9^ uncovers the structural basis of selectivity for lasmiditan (Fig. 1f-h). Comparison of 5-HT_1F_ with 5-HT_1B_ shows that the ligand-receptor interaction are basically conserved in OBP, while differences in EBP, which are formed by TM4/5/6, and ECL2. For the TM4/5, the residues that interact with the trifluorobenzene group of lasmiditan in 5-HT_1F_ are highly conserved with 5-HT_1B_ (Fig. 1f). However, the conformation of TM4/5 shows significant changes between 5-HT_1F_ and 5-HT_1B_. On the extracellular side, TM4 shifts 3.6 Å, and TM5 shifts 2.8 Å for 5-HT_1F_ from those in 5-HT_1B_ (Fig. 1f). For the TM6, the E^6×55^ and N^6×59^ of 5-HT_1F_ form hydrogen bonds with lasmiditan, but the corresponding residues are S^6×55^ and P^6×59^ in 5-HT_1B_, and they cannot provide the corresponding interactions. For the ECL2, including the region that interacts with lasmiditan shows different conformations between 5-HT_1F_ and 5-HT_1B_ (Fig. 1f).

5-HT_1E_ is the receptor with the highest homology to 5-HT_1F_. The structure of 5-HT_1E_ we previously reported reveals the mechanism of the 5-HT_1E/1F_ selective ligand BRL54443 for 5-HT_1E_. Lasmiditan is only selective for 5-HT_1F_, rather than for 5-HT_1E_. Structural comparison of 5-HT_1F_ and 5-HT_1E_ provides an opportunity to uncover the mechanism of selectivity of lasmiditan on 5-HT_1F_. Although 5-HT_1F_ and 5-HT_1E_ have relatively conserved residues for ligand binding both in OBP and EBP, the conformations of TM4, TM5, and ECL2 show significant differences. Among them, the extracellular end of TM4 shifted by 3.5 Å, TM5 shifted by 2.0 Å, and the backbone of ECL2 shows different conformarion (Fig. 1g). These changes are similar in comparison with 5-HT_1B_ (Fig. 1f). We further compared the 5-HT_1F_ structure with other 5-HT_1_ subfamily receptors, the results showed that the conformation of the TM4-TM5-ECL2 region is relatively conserved in 5-HT_1A_, 5-HT_1B_, 5-HT_1D_, and 5-HT_1E_, but not for 5-HT_1F_ (Fig. 1h; Supplementary information, Fig. S3b-d). To confirm the roles of ECL2 on ligand selectivity, we replaced the ECL2 of 5-HT_1F_ with that of other 5-HT_1_ receptors and tested the receptor activation. The result shows that the lasmiditan induced activation was significantly affected (Supplementary information, Fig. S3e). The importance of EPB for lasmiditan binding and the different shape of EBP of 5-HT_1F_ from other 5-HT receptors determines the high selectivity of lasmiditan on 5-HT_1F_.

The lasmiditan induces activation for 5-HT_1F_ undergoing a canonical conformational rearrangement. Comparing to the inactive state 5-HT_1B_^10^, lasmiditan triggers the toggle switch residue W^6×48^ of 5-HT_1F_ downward movement, then induces the conformational changes in PIF, DRY, and NPxxY motifs (Supplementary information, Fig. S4a-e).These conformational changes further cause an 8 Å outward movement of TM6, which allows the α5 helix of Gα_i1_ insert into the intracellular cavity formed by the receptor TMD bundle, a hallmark of GPCR activation (Supplementary information, Fig. S4f). Structural comparison of the G_i_-coupled 5-HT_1F_ with the G_o_-coupled 5-HT_1B_ complexes^9^ reveals differences in G-protein coupling (Supplementary information, Fig. S4g, h). Although the conformation of the main interfaces between receptor and G-protein are similar, the G-protein conformation shows observable changes. The main body of the Ras-like domain shares a similar conformation, while the N-terminus of αN shift 9.4 Å and the last residue of α5 shift 2.4 Å between G_i_ and G_o_ (Supplementary information, Fig. S4g). Comparing the 5-HT_1F_–G_i_ complex with other 5-HT_1_–G_i/o_ complex structures^7,9^, we found that the αN of 5-HT_1F_ bound G_i_ shifts away from other 5-HT_1_ receptors bound G_i/o_, which suggests that the coupling of 5-HT_1F_ to G_i_ protein is unique from other 5-HT_1_ receptors (Fig. 1i).

In summary, in this paper, we report the cryo-EM structure of the 5-HT_1F_–Gi complex bound to a highly selective anti-migraine drug lasmiditan. The structure reveals the binding mode of lasmiditan in 5-HT_1F_. Comparison of our structure with the previously reported 5-HT_1_ structures^9^ provide the basis of the selectivity for lasmiditan to 5-HT_1F_. The determination for selectivity mainly attributes to the interaction between the trifluorobenzene group of lasmiditan and the specific extended binding pocket (EBP) of 5-HT_1F_. Furthermore, our structure reveals a conserved mechanism for activation of 5-HT_1F_ and the unique G protein coupling conformation from that in the 5-HT_1_–G-protein structures^7,9^. Together, these results provide a rational template for design of new generation of anti-migraine drugs that selectively target 5-HT_1F_, therefore avoiding the main disadvantage of cardiovascular side effects associated with the triptan class of anti-migraine drugs.

## Data availability

The corresponding coordinates and cryo-EM density map have been deposited in the Protein Data Bank (http://www.rcsb.org/pdb) with code 7EXD, and in EMDB (http://www.ebi.ac.uk/pdbe/emdb/) with code EMD-31371.

## Acknowledgements

The cryo-EM data were collected at the Cryo-Electron Microscopy Research Center, Shanghai Institute of Materia Medica, Chinese Academy of Sciences (Shanghai, China). This work was partially supported by the Ministry of Science and Technology (China) grant (2018YFA0507002 to H.E.X.); CAS Strategic Priority Research Program (XDB37030103 to H.E.X.); Shanghai Municipal Science and Technology Major Project (2019SHZDZX02 to H.E.X.); the National Natural Science Foundation (31770796 to Y.J.); National Science & Technology Major Project “Key New Drug Creation and Manufacturing Program” (2018ZX09711002-002-002 to Y.J.).

## Author contributions

S.H. and P.X. designed the expression constructs, purified the complexes, prepared samples for negative stain and data collection toward the structures, performed functional assay, prepared the figures and manuscript draft. P.X. evaluated the specimen by negative-stain EM, screened the cryo-EM conditions, prepared the cryo-EM grids, collected cryo-EM images, built the model, and refined the structures. Y.T., C.Y., and Y.Z. participated in the NanoBiT G-protein recruitment assays. Y.J. participated in the supervision of S.H., P.X., Y.T., C.Y., and Y.Z. and analyzed the structures, edited the manuscript. H.E.X. conceived and supervised the project, analyzed the structures, and wrote the manuscript with inputs from all authors.

## Supplementary information

### Supplementary information, Figure S1

**Fig. S1.**
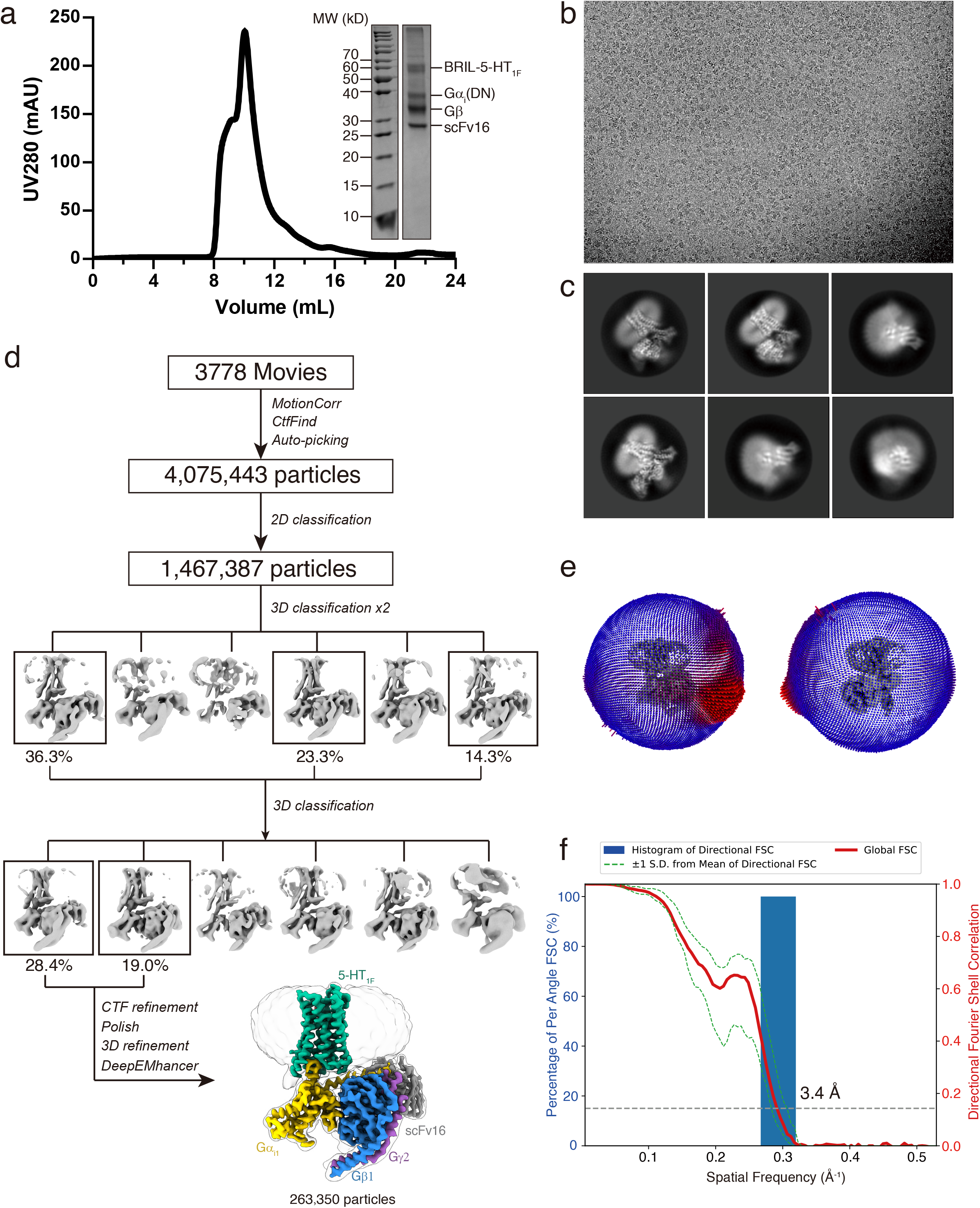
Sample preparation and cryo-EM of the 5-HT_1F_–G_i1_ complexes. **a** Analytical size-exclusion chromatography of the purified complex and SDS-PAGE/Coomassie blue stain of the purified complex. **b, c** Representative cryo-EM image and 2D averages. **d** Flowchart of cryo-EM data analysis of the lasmiditan bound-5-HT_1F_–G_i_ complex. **e** Euler angle distribution of particles used in the final reconstruction. **f** ‘Gold-standard’ Fourier shell correlation curves of the lasmiditan–5-HT_1F_–G_i_ complex.

### Supplementary information, Figure S2

**Fig. S2.**
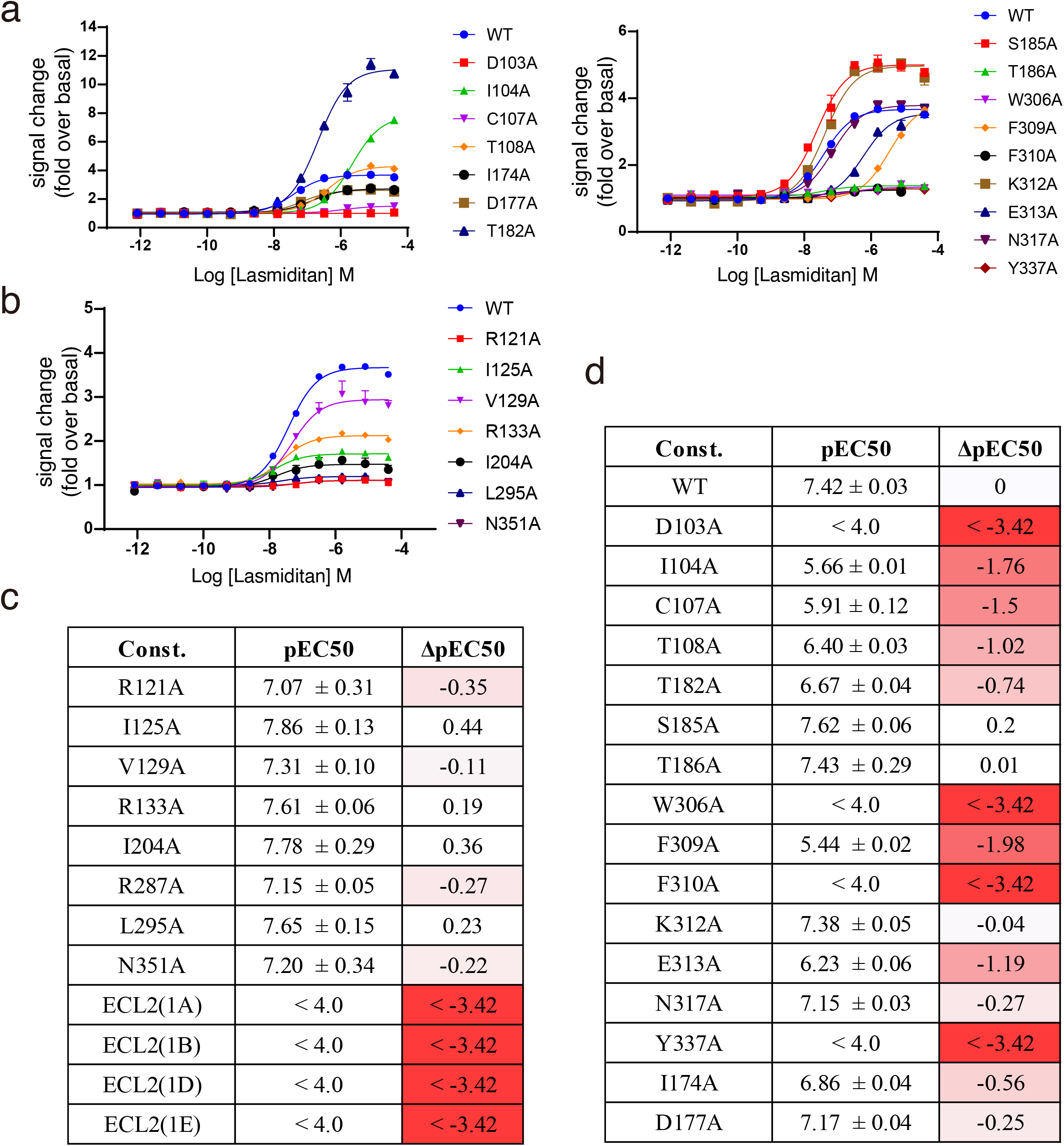
Mutagenesis data of lasmiditan mediate 5-HT_1F_ activation by NanoBiT Gi-protein-recruitment assay. **a** Dose response cuves of mutations on ligand-binding pocket. **b** Dose response cuves of mutations on G-protein interaction interface. **c** pEC_50_ of mutations on G-protein interaction interface and ECL2 chimeraic receptors. **d** pEC_50_ of mutations on ligand-binding pocket. Data are mean ± s.e.m. from at least three independent experiments performed in technical triplicate.

### Supplementary information, Figure S3

**Fig. S3.**
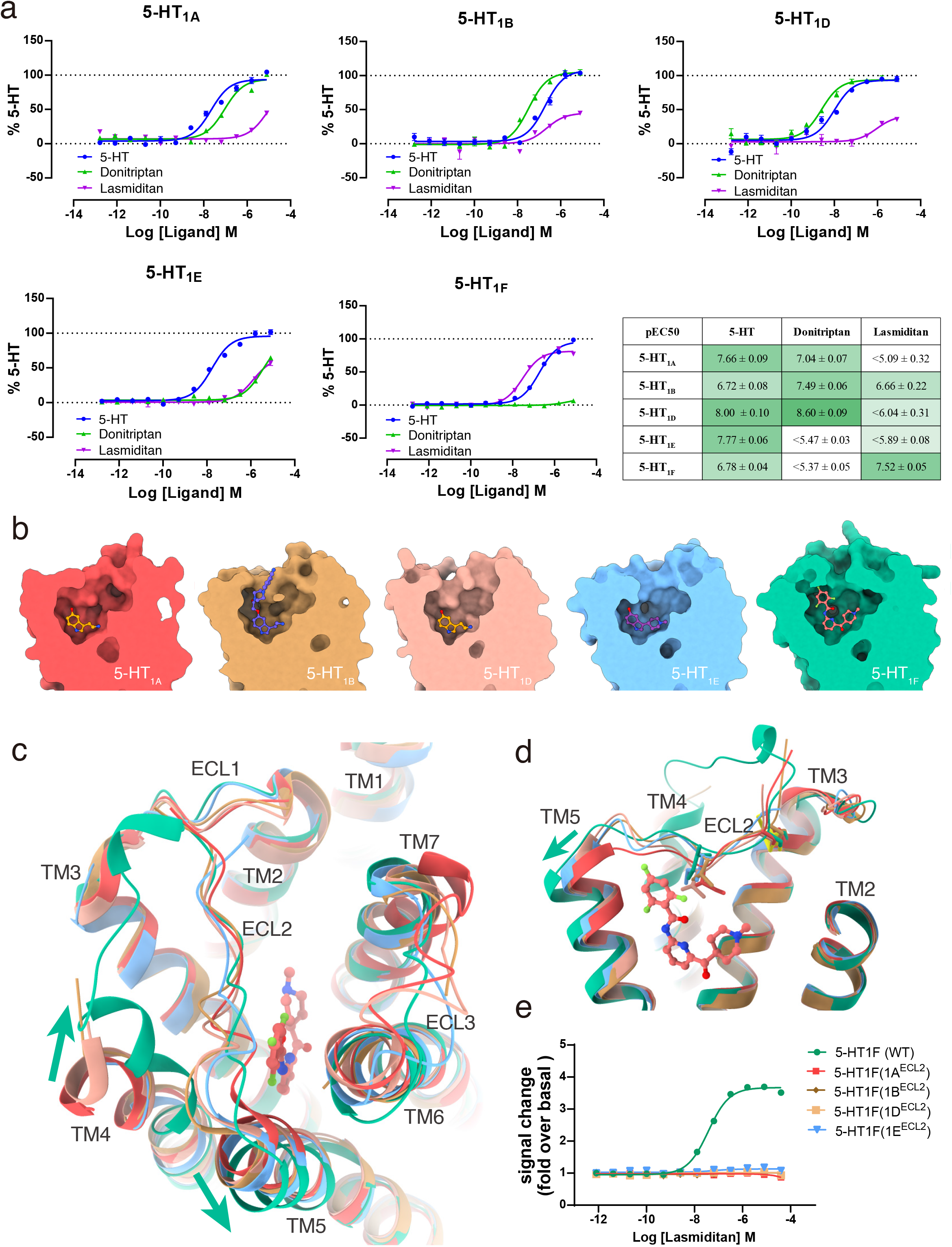
Selectivity of lasmiditan. **a** Dose response cuves and pEC50 of 5-HT, donitriptan, and lasmiditan induced activation of 5-HT_1A_, 5-HT_1B_, 5-HT_1D_, 5-HT_1E_ and 5-HT_1F_. b comparison of the ligand-binding pocket among the structures of G_i/o_ coupled 5-HT_1A_ (red, PDB code: 7E2Y), 5-HT_1B_ (tan, PDB code: 6G79), 5-HT_1D_ (yellow, PDB code: 7E32), 5-HT_1E_ (blue, PDB code: 7E33) and 5-HT_1F_ (green). **c, d** Comparison of TM4-TM5-ECL2 region among 5-HT_1_ sub-family receptors. c, extracellular view; d, side view. **e** The replacement of ECL2 of 5-HT_1F_ with other 5-HT_1_ receptors affects the lasmiditan mediate activation. Data are mean ± s.e.m. from at least three independent experiments performed in technical triplicate.

### Supplementary information, Figure S4

**Fig. S4.**
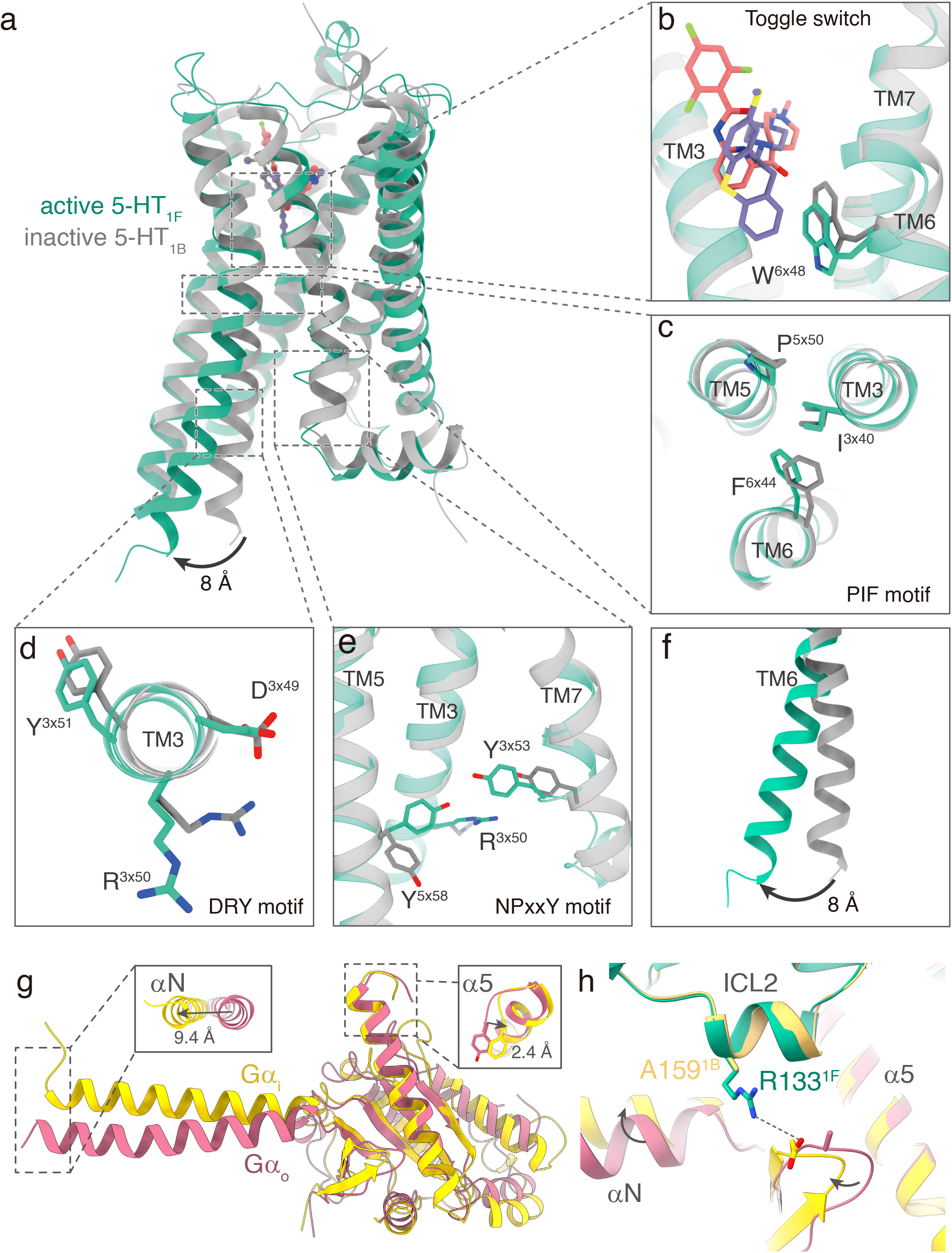
Lasmiditan induced activation of 5-HT_1F_. **a** Structural uperposition of active 5-HT_1F_ and inactive 5-HT_1B_ (PDB code: 5v54) complexes. **b-e** residues rearrangement of Toggle switch (**b**), PIF motif (**c**), DRY motif (**d**), and NPxxY motif (**e**). **f** The outwad movement of intracellular end of TM6. **g** Comparison of the 5-HT_1F_ bound Gα_i_ conformation and 5-HT_1B_ bound Gα_o_ conformation. The left arro indicates the movement of αN helix in Gα_i_ structure relative to Gα_o_ structure. The right arrow indicates the movement of the last amino acid in the Gα α5 helix in Gα_i_ structure relative to Gα_o_ structure. **i** Comparison of ICL2-G protein interactions in the 5-HT_1F_–G_i_ and 5-HT_1B_–mG_o_ structures.

## Materials and methods

### Constructs of 5-HT_1F_ and G_i1_ heterotrimer

The full-length gene sequences of wild type *human* 5-HT_1F_ receptors were subcloned into pFastbac vector using ClonExpress II One Step Cloning Kit (Vazyme Biotech Co.,Ltd). An N-terminal thermally stabilized BRIL^1^ as a fusion protein to enhance receptor expression. N-terminal fusions of Flag tag and 8×His tag were used to facilitate protein purification. A dominant-negative Gα_i1_ was generated by site-directed mutagenesis to incorporate mutations S47N, G203A, A326S, and E245A that improves the dominant-negative effect by weakening a salt bridge that helps to stabilize the interactions with the βγ subunits^2^. All the three G_i_ subunits, *human* DN_Gα_i1_, wild type Gβ1, and Gγ2 were cloned into the pFastBac vector separately. A no-tag single-chain antibody scFv16^3^ was cloned into pFastBac.

### Insect cell expression

*Human* 5-HT_1F_, DNGα_i1_, Gβ_1_, Gγ_2_, and scFv16 were co-expressed in *Trichoplusia ni* Hi5 insect cells using the baculovirus method (Expression Systems). Cell cultures were grown in ESF 921 serum-free medium (Expression Systems) to a density of 2-3 million cells per ml and then infected with four separate baculoviruses at a suitable ratio. The culture was collected by centrifugation 48 h after infection and cell pellets were stored at −80°C.

### Complex purification

The complex was purified as previously described^4^. In brief, cell pellets were thawed in 20 mM HEPES pH 7.4, 50 mM NaCl, 10 mM MgCl_2_ supplemented with Protease Inhibitor Cocktail (Bimake). The 5-HT_1F_ complex formation was initiated by addition of 10 μM lasmiditan (TargetMol), apyrase (25 mU/ml, Sigma). The suspension was incubated for 1 h at room temperature and the complex was solubilized from the membrane using 0.5% (w/v) n-dodecyl-β-d-maltoside (DDM, Anatrace) and 0.1% (w/v) cholesteryl hemisuccinate (CHS, Anatrace) for 2 h at 4°C. Insoluble material was removed by centrifugation at 65,000 g for 30 min and the solubilized complex was immobilized by batch binding to Talon affinity resin. The resin was then packed and washed with 20 column volumes of 20 mM HEPES pH 7.4, 100 mM NaCl, 5 mM MgCl_2_, 0.01% (w/v) LMNG, and 0.002% (w/v) CHS, 10 μM lasmiditan. Finally, the complex was eluted in buffer containing 300 mM imidazole and concentrated with an Amicon Ultra Centrifugal Filter (MWCO 100 kDa). Complex was subjected to size-exclusion chromatography on a Superdex 200 Increase 10/300 column (GE Healthcare) pre-equilibrated with 20 mM HEPES pH 7.4, 100 mM NaCl, 0.05% (w/v) digitonin, and 10 μM lasmiditan, to separate complex from contaminants. Eluted fractions consisting of receptor and G_i_-protein complex were pooled and concentrated.

### NanoBiT G-protein recruitment assay

Analysis of G-protein recruitment was performed by using a modified protocol of NanoBiT system (Promega) assay described previously^5^. Receptor-LgBiT, Gα_i1_, SmBiT-fused Gβ_1_, and Gγ_2_ were co-expressed in *Trichoplusia ni* Hi5 insect cells using the baculovirus method (Expression Systems). Cell cultures were grown in ESF 921 serum-free medium (Expression Systems) to a density of 2-3 million cells per ml and then infected with four separate baculoviruses at a suitable ratio. The culture was collected by centrifugation 48 h after infection and cell pellets were collected with PBS. The cell suspension was dispensed in a white 384-well plate at a volume of 40 μl per well and loaded with 5 μl of 90 μM coelenterazine diluted in the assay buffer. Test compounds (5 μl) were added and incubated for 3-5 min at room temperature before measurement. Luminescence counts were normalized to the initial count and fold-change signals over vehicle treatment were used to show G-protein binding response.

### Cryo-EM grid preparation and data collection

For the preparation of cryo-EM grids, 3 μL of the purified lasmiditan–5-HT_1F_–G_i_ complex at concentration ∼15 mg/ml were applied onto a glow-discharged holey carbon grid (Quantifoil R1.2/1.3 Au 300). Grids were plunge-frozen in liquid ethane using Vitrobot Mark IV (Thermo Fisher Scientific). Frozen grids were transferred to liquid nitrogen and stored for data acquisition. Cryo-EM imaging was performed on a Titan Krios at 300 kV using Gatan K3 Summit detector in the Cryo-Electron Microscopy Research Center, Shanghai Institute of Materia Medica, Chinese Academy of Sciences (Shanghai, China). The images were recorded at a dose rate of about 26.7 e^-^/Å^2^/s with a defocus ranging from −1.2 to −2.2 μm. The total exposure time was 3 s and intermediate frames were recorded in 0.083 s intervals, resulting in a total of 36 frames per micrograph.

### Image processing and map construction

Dose-fractionated image stacks were aligned using MotionCor2.1^6^. Contrast transfer function (CTF) parameters for each micrograph were estimated by Gctf^7^. Cryo-EM data processing was performed using RELION-3.1^8^. Automated particle picking yielded 4,075,443 particles that were subjected to reference-free 2D classification to discard poorly defined particles, producing 1,467,387 particles. The map of 5-HT_1B_–miniG_o_ complex (EMDB-4358)^9^ low-pass filtered to 60 Å was used as an initial reference model for 3 rounds of 3D classification. Two subsets show the high-quality receptor density was selected, producing 263,350 particles. The selected subsets was subsequently subjected to 3D refinement, CTF refinement, Bayesian polishing, and DeepEMhancer^10^. The final refinement generated a map with an indicated global resolution of 3.4 Å at a Fourier shell correlation of 0.143. Local resolution was determined using the 3DFSC package with half maps as input maps.

### Model building and refinement

The cryo-EM structure of 5-HT_1B_–mG_o_ complex (PDB code: 6G79) and the G_i_ protein model (PDB code: 6PT0) were used as the start for model rebuilding and refinement against the electron microscopy map. The model was docked into the electron microscopy density map using Chimera^11^, followed by iterative manual adjustment and rebuilding in COOT^12^ and ISOLDE^13^. Real space and reciprocal space refinements were performed using Phenix programs^14^. The model statistics were validated using MolProbity^15^. Structural figures were prepared in Chimerax^16^.

